# Mediator complex subunit Med19 binds directly GATA DNA-binding zinc finger and functions with Med1 in GATA-driven gene regulation *in vivo*

**DOI:** 10.1101/2020.04.03.023895

**Authors:** Clément Immarigeon, Sandra Bernat-Fabre, Emmanuelle Guillou, Alexis Verger, Elodie Prince, Mohamed A. Benmedjahed, Adeline Payet, Marie Couralet, Didier Monte, Vincent Villeret, Henri-Marc Bourbon, Muriel Boube

## Abstract

The evolutionarily-conserved multiprotein Mediator complex (MED) serves as an interface between DNA-bound transcription factors (TFs) and the RNA Polymerase II machinery. It has been proposed that each TF interacts with a dedicated MED subunit to induce specific transcriptional responses. However, binary MED subunit - TF partnerships are probably oversimplified models. Using *Drosophila* TFs of the GATA family - Pannier (Pnr) and Serpent (Srp) - as a model, we have previously established GATA cofactor evolutionarily-conserved function for the Med1 Mediator subunit. Here, we show that another subunit, Med19, is required for GATA-dependent gene expression and interacts physically with Pnr and Srp *in cellulo, in vivo* and *in vitro* through their conserved C-zinc finger (ZF), indicating general GATA co-activator functions. Interestingly, Med19 is critical for the regulation of all tested GATA target genes which is not the case for Med1, suggesting differential use of MED subunits by GATAs depending on the target gene. Lastly, despite their presumed distant position within the MED middle module, both subunits interact physically. In conclusion, our data shed new light first on the MED complex, engaging several subunits to mediate TF-driven transcriptional responses and second, on GATA TFs, showing that ZF DNA-binding domain also serves for transactivation.

## Introduction

Transcription, the first stage of gene expression, is a fundamental cellular process governed by the binding of sequence-specific transcription factors (TFs) at gene enhancers, inducing the recruitment/activation of the general RNA Polymerase II (Pol II) machinery at gene promoters. In eukaryotes, TFs do not bind directly the Pol II enzyme but instead contact a multisubunit complex called Mediator (MED), serving as a physical and functional interface between DNA-bound TFs and PolII (for review see (1–3)). Whereas TF DNA-binding specificity has been largely decoded, how TFs interact with the Mediator complex has been less extensively studied, and it is not clear whether each TF binds a specific MED subunit or whether TF-MED interactions obey more complex rules.

Mediator is an evolutionarily-conserved complex composed of 25 to 30 distinct proteins distributed in four modules: head, middle, and tail forming the core MED, and a separable regulatory Cdk8 kinase module (CKM) (1). Despite a general role of the Mediator complex in regulating transcription, some MED subunits display striking functional specificities, as exemplified by their differential requirements for cell viability (4, 5), their involvement in specific human diseases (6, 7), or their roles in given developmental processes (8–10). It has been proposed that MED subunit specificity comes from their ability to contact specific transcription factors and mediate their regulatory activity (11, 12). For example, specific interactions and cofactor activities have been demonstrated between Med15 and SMAD transcription factors in Xenopus (13), Med23 and RUNX2 in mice (14) Med12 and Gli3 in mammalian cells (15), Med19 and REST in mammals or Med19 and HOX developmental regulators in *Drosophila* (16, 17), or also between Med1 and hormone nuclear receptors or GATA TF families in mammalian cells (18, 19).

Mammalian GATA TF family comprises 6 members (GATA1-6), with conserved homologs among both vertebrates and invertebrates (20). They generally contain two highly-conserved zinc finger (ZF) domains, of which the C-terminal one (C-ZF) is both necessary and sufficient for sequence-specific DNA binding at WGATAR genomic sites, while the N-terminal ZF (N-ZF) appears only to modulate DNA-binding affinity (21) and has been involved in direct interactions with GATA cofactors (22–25). Mammalian GATAs are key regulators of developmental processes: GATA1, -2 and -3 are crucial hematopoietic TFs while GATA4, -5 and -6 control cardiac development, among other functions (20). Interestingly, among the 5 GATA TFs encoded by the *Drosophila* genome, only Serpent (Srp), is a *bona fide* hematopoietic GATA factor, while another one, Pannier (Pnr), is involved in cardiac development (26). GATA/Pnr activity is also crucial during central thorax patterning and dorso-central (DC) mechanosensory bristle formation, and it has been studied in depth in this context (27–29)(28). Within the wing imaginal disc, the Pnr TF directly activates proneural genes of the *achaete-scute* (*ac-sc*) complex in the dorso-central cluster, which gives rise to the DC bristles (27). In addition, Pnr activates *wingless* in a strip of cells of the presumptive centro-lateral notum (30).

A genome-wide RNA interference screen in *Drosophila* cultured cells allowed us identifying a set of MED subunits as modulators of GATA/Serpent-induced transactivation, among which, Med12, Med13, Med1 and Med19 (31). This work further showed that Med12 and Med13 subunits are required *in vivo* for Srp-driven developmental processes, but we were unable to detect direct physical interaction with Srp *in vitro*, suggesting that GATA/Srp may recruit the Mediator complex by contacting other MED subunits. To address the question of evolutionary conservation of MED subunits-TFs partnerships, we recently asked whether the Med1 subunit mediates also GATA TFs function in *Drosophila* (32). We showed that *Drosophila* Med1 does interact physically with both Pnr and Srp GATA TFs, through their conserved zinc finger region. Furthermore, *in vivo* experiments showed that Med1 is involved in Srp-driven hematopoiesis and Pnr-driven thorax differentiation and is required for Srp and Pnr target gene expression in the corresponding tissues. These data both established that the Med1 GATA cofactor is evolutionarily conserved and involves the GATA N- and C-zinc finger domains. Nevertheless, we also showed that *Drosophila* Med1 is critical for *achaete*-but not for *wingless*-induced transactivation by Pnr, raising the possibility that other MED subunits could mediate GATA TFs functions.

Here, we reveal that another MED subunit, Med19, also acts as a GATA coactivator. *Med19* mutants phenocopy *pnr* loss-of-function, and extinguish the expression of both Pnr target genes *achaete* and *wg*. Using Immunoprecipitation, pull-down and Bimolecular Fluorescence Complementation (BiFC) techniques, we establish that Med19 physically interacts with Pnr *in cellulo, in vivo* and *in vitro* through its C-ZF domain. We further show that Med19 also interacts physically with Srp, suggesting that, like Med1, Med19 acts as a general GATA cofactor. Moreover, we show that both Med1 and Med19 jointly regulate a series of Srp target genes in *Drosophila* cultured cells. Finally, *in vitro* experiments revealed that Med1 and Med19 physically interact through the Med1 domain conserved throughout eukaryotes. Taken together, our results show that GATA-driven regulatory functions in *Drosophila* require the close collaboration of two MED complex subunits. The evolutionary conservation of Med19 and GATA interacting domains suggests that Med19 may play a conserved GATA cofactor function in mammals.

## Results

### *Drosophila* Med19 is required for notum morphogenesis, bristle development and GATA/Pannier target gene activation

Our study of the first-characterised metazoan mutant for *Med19* revealed a Med19 function as a specific co-activator of HOX TF family during *Drosophila* development (16). Our whole-genome dsRNA screen in *Drosophila* cultured cells previously identified Med19 as one of several MED subunits capable of modulating Srp TF-induced transactivation. (31). This led us to ask whether and how Med19 could interact with GATA TFs *in vivo*. To this end, we generated *Med19* mutant clones in the larval wing imaginal disc, giving rise to adult thoracic structures whose proper development depends on GATA/Pnr activity. Flies bearing *Med19*^*-*^ clones displayed specific phenotypes in the thorax, including thoracic cleft and loss of dorso-central (DC) mechanosensory macrochaetes (Fig.1A, D), typical of *pnr* loss-of-function (28, 29). We observed similar phenotypes upon expression of interfering RNAs (RNAi) against *Med19* in the *apterous (ap)* domain encompassing all the presumptive notum (Fig S1A-B). To investigate functional relationship between Med19 and Pnr, we first examined *pnr* gene expression in *Med19*-deficient wing discs by fluorescent *in situ* hybridization (FISH), and observed that *pnr* is expressed in Med19-depleted wing discs (Fig S1C-F). Thus, *Med19* mutant phenotypes cannot be explained by a loss of *pnr* expression. To further investigate the functional relationship between Med19 and Pnr, we then examined GATA/Pnr TF activity in *Med19* loss-of-function clones by analyzing the expression of known Pnr target genes. Compared to wild-type cells shown in Fig. 1B-C”, we observed that both *wingless (*wg) and *achaete (ac)* expression was cell autonomously lost in *Med19*^*-/-*^ cells (Fig.1E-F”) indicating that Med19 is required for Pnr target gene expression. Note that *ac* expression has been visualized by a DC-*ac-*lacZ reporter gene which is directly activated upon Pnr binding to the DC *ac* enhancer (27).

**Figure 1.**
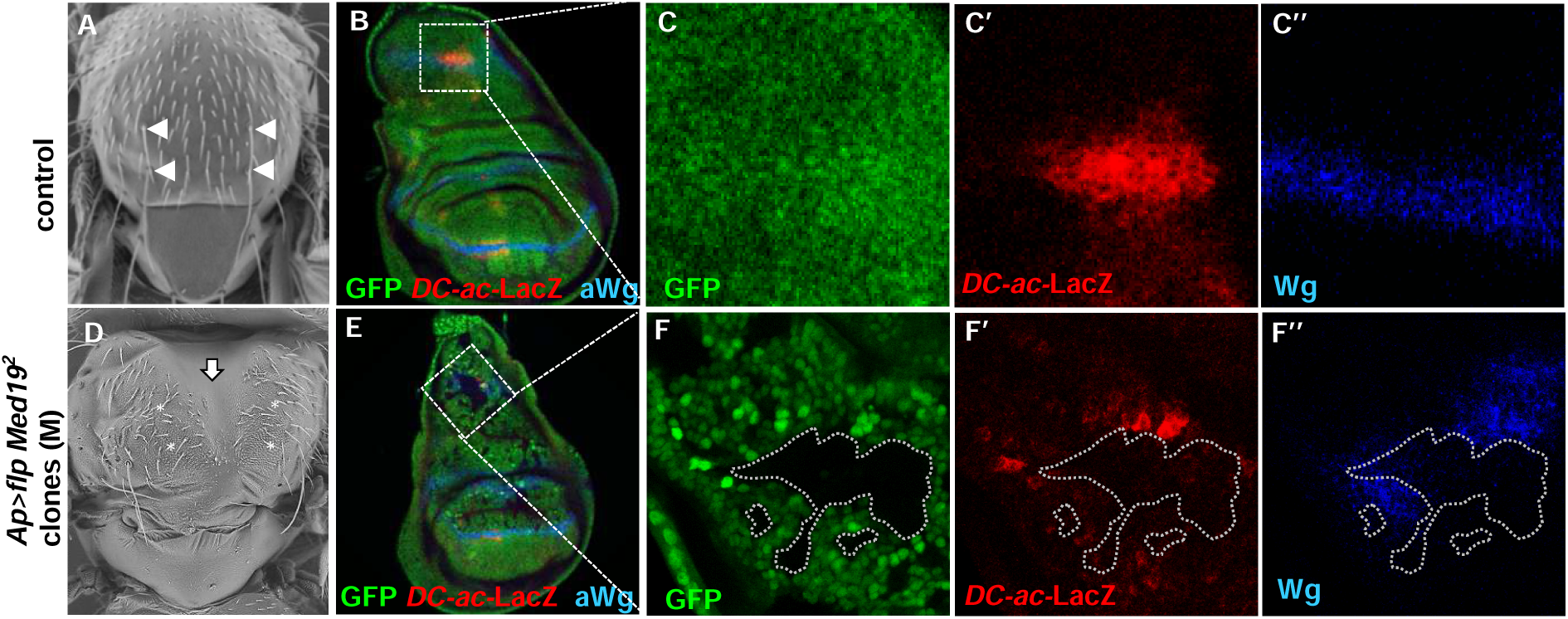
*Med19* is required for GATA Pannier target gene activation. *Drosophila* thorax: (**A**) wild type or (**D**) bearing *Med19*^*-/-*^ clones. Arrows point to DC bristles in A and asterisks to their expected position in D. The large vertical arrow points to the thoracic cleft. Immunofluorescence analysis of Pannier target genes expression in control wing discs (**B-C**) and in *Med19*^*-/-*^ mitotic clones (GFP-) generated in the dorsal compartment giving rise to the presumptive thorax and wings) (**E-F**). The expression of *DC-ac-*LacZ reporter and Wg protein are revealed with anti-βgal (red), and anti-Wg (blue) antibodies. Magnifications of the DC region are shown (C and F).

These data show that Med19 is cell autonomously required for Pnr activity but not for Pnr expression, suggesting that it could act as a GATA/Pnr cofactor.

### *Drosophila* Med19 interacts physically with the Pannier GATA TF

We investigated whether GATA/Pnr transcription factor and Med19 physically interact by using three independent experimental approaches: co-immunoprecipitation (co-IP) from cultured cells, *in vitro* pull-down and *in vivo* Bimolecular Fluorescence Complementation (BiFC) interaction tests. We first tested whether Pnr-MED complexes actually form within *Drosophila* cells by performing co-Immunoprecipitations (co-IP) experiments on total protein extracts from cultured cells expressing a functional Myc-tagged Pnr form. We observed that Pnr co-precipitated with endogenous *Drosophila* Med19 (Fig.2A). In the reverse experiment, endogenous Med19 protein co-precipitated with Myc-tagged Pnr protein (Fig.2B). Altogether, these data provide complementary evidence for the formation of Med19-GATA complexes in *Drosophila* cells.

**Fig. 2:**
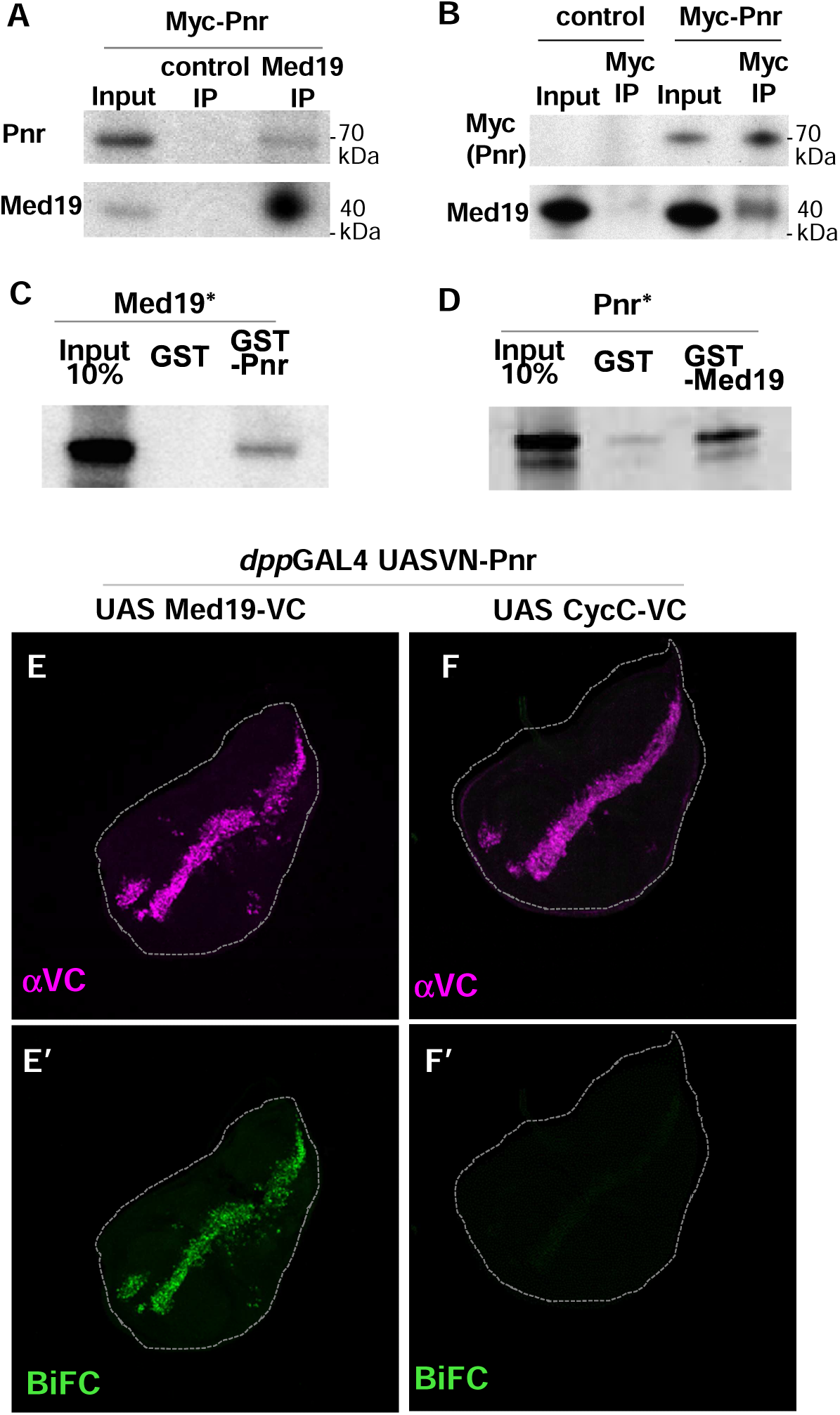
Med19 physically interacts with GATA/Pnr *in cellulo, in vitro* and *in vivo.* Co-immunoprecipitation (IP) experiments from S2 cells transfected with *pAct*-Myc-Pnr using anti Med19 antibody or preimmune serum (control IP) **(A**). The reverse experiment was performed using anti-Myc beads from control cells or cells expressing Myc-Pnr (**B**). Western blot assays using αMed19, αPnr or αMyc antibodies are shown. Autoradiographs from GST pull-down assays between GST-Pnr and ^35^S-labeled (*) *in vitro*-translated Med19 (**C**), and between GST-Med19 and *in vitro*-translated ^35^S-Pnr (**D**).

To investigate whether Med19-Pnr interaction is direct, we tested the ability of Med19 and Pnr proteins to bind each other physically *in vitro* through pulldown with Glutathione S-transferase (GST) fusion proteins. *In vitro*-produced Med19 readily bound full-length recombinant GST-Pnr (Fig.2C), and *vice versa* (Fig.2D). These results show that Med19 and Pnr can interact physically in the absence of any other *Drosophila* factor, especially other MED subunits.

We then used Bimolecular Fluorescence complementation (BiFC) technique (16, 33) to tackle the question of Med19-GATA molecular interaction *in vivo*. Based on fusing N- and C-terminal portions (VN and VC) of the GFP-variant Venus protein, with two proteins of interest respectively, this technique allows the reconstitution of a fluorescent Venus protein if the two candidate proteins are close enough within the cell. We used the *dpp*GAL4 driver (Gal4/UAS system (34)) to co-express VN-Pnr with Med19-VC or another MED subunit fusion, CycC-VC, at the posterior (A/P) frontier of the wing imaginal disc (Fig2.E, F). The co-expression of VN-Pnr and Med19-VC resulted in a clear BiFC signal in the *dpp*GAL4 expression domain, whereas the control VN-Pnr /CycC-VC combination gave a very low signal (Fig.2E’-F’), even though CycC-VC and Med19-VC fusion proteins were expressed at similar levels (Fig. 2E-F). These data indicate that the BiFC technique discriminates specific interactions between different subunits within the MED complex and show that Med19 and Pnr are in close proximity in the nucleus of living cells. Of note, the BiFC signal was observed in the entire *dpp*GAL4 expression domain including the wing pouch where GATA/Pnr is not normally expressed, showing that the Pnr-Med19 interaction can occur at ectopic locations independently of Pnr endogenous partners, thus providing further support for a direct molecular interaction *in vivo*.

Collectively, *in cellulo, in vitro* and *in vivo* data support a direct physical interaction between the GATA/Pnr transcription factor and the Med19 Mediator subunit and suggest that the Pnr TF functionally interacts with the entire MED complex via a direct molecular contact with its Med19 subunit.

### Med19 Core and HIM domains bind the C-zinc finger domain of GATA/Pnr

We previously showed that Med1 directly interacts with the dual Zinc Finger domains of Pnr and that HOX TFs bind so-called Med19 HIM (Homeodomain-interacting motif) domain. We therefore decided to further investigate two aspects: whether or not Med19 and Med1 interact with the same Pnr domain and whether or not Med19 binds GATA and HOX TFs through the same domain.

We first looked for the Med19–interacting domain(s) within the GATA/Pnr protein using full length GST-Med19 as a bait (Fig.3A). Pnr was split into three parts: the poorly evolutionarily-conserved N-terminal region (amino acids (aa) 1-137), the strongly-conserved central region spanning the 2 zinc fingers, N- and C-ZF, and the divergent C-terminal region containing two amphipathic alpha helices, H1 and H2. Only the ZF-containing region (aa 130-278) displayed significant binding. When cutting full-length Pnr into two halves separating the two zinc finger domains, binding was observed only with the C-ZF-containing part (Fig. 3A), suggesting that C-ZF mediates binding of Pnr to Med19. Consistently, the ability of the N-ZF-containing half of Pnr to bind Med19 was recovered when we added back the C-ZF proper (aa 220-253) containing the four zinc-chelating cysteines forming the finger structure. Interestingly, binding was increased when the C-ZF proper was extended by its neighboring C-terminal 25 amino acids (basic tail motif, aa 253-278; Fig. 3B). Sequence alignment of *Drosophila* and mammalian GATA C-ZF domains indicates that the basic tail region has been strongly conserved during evolution, especially at positions which have been shown to participate in DNA binding (open circles in Fig. 3B, (35, 36)). Together, these experiments indicate that within the Pnr C-ZF domain, the proper zinc-finger subdomain and its adjacent basic tail are both necessary for optimal Med19 binding.

**Fig. 3:**
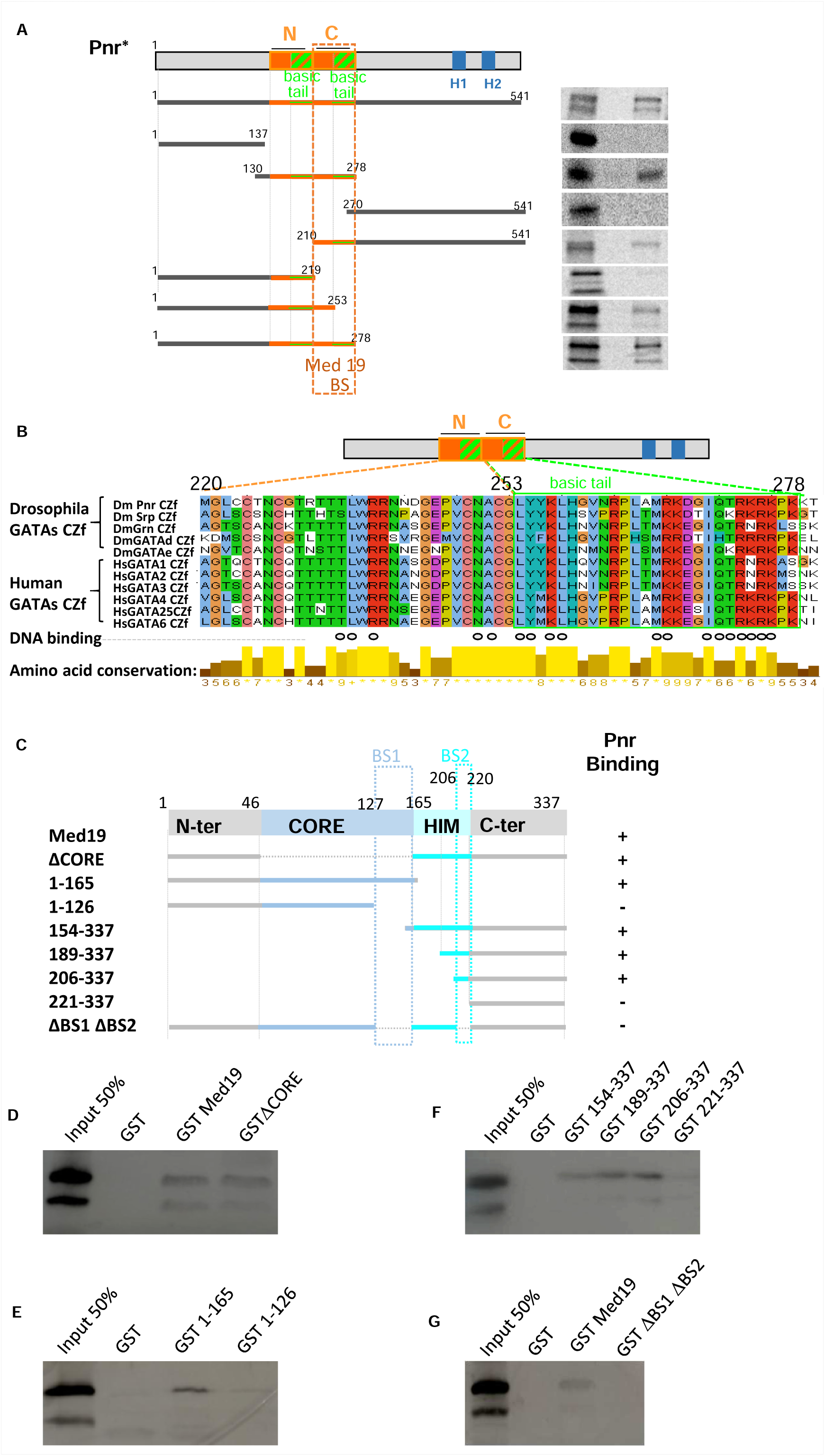
GATA/Pnr C-zinc finger domain interacts with two conserved Med19 domains. GST pulldown assays allows delimitating interacting domains within Med19 and Pnr proteins. Schematic representation of GATA-Pnr and the multiple Pnr fragments generated to probe for binding to full-length GST-Med19 (**A**). N and C show proper zinc fingers (orange boxes) and their respective basic tails (hatched green and orange boxes). H1 and H2 show amphipathic alpha helices. Autoradiographs from GST pulldown experiments are shown on the right side for each fragment. The critical domain for strong binding is narrowed down to Pnr amino-acids 220-278. This domain comprising C-ZF proper zinc finger and its basic tail is highly conserved in *Drosophila* GATA factors (Srp, Grn, GATAd and GATAe) as well as human GATA factors (GATA1 to 6) as shown in panel **B**. Open circles denote residues participating in DNA binding (34, 35). Level of amino acid conservation is represented in beneath. Schematic representation of Med19 and the multiple Med19 fragments generated as GST fusions (**C**). For each truncated construct, the solid bar represents the portion of Med19 that is included. Numbers correspond to amino-acids. + and – summarize experimental results for Pnr binding, deduced from HA-Pnr 1-291 detections after GST-pulldowns (Western blots are shown in panels D to G). 2 distinct domains of Med19 are sufficient for binding Pnr (BS1 and BS2 in A).

In the reciprocal experiment, we examined the GATA/Pnr interacting domain within the Med19 protein. Our prior analysis of *Drosophila* Med19 function and evolutionary conservation within the eukaryotic kingdom (16, 37), allowed to define four structural domains: a conserved MED-anchoring “CORE” region, an animal-specific basic HOX homeodomain-interacting motif (HIM) and two less well conserved N-and C-terminal regions. To investigate which protein domain(s) is (are) required for Pnr binding, we tested the ability of *in vitro* translated Pnr 1-291 HA tagged form to bind a series of GST-Med19 truncated forms (Fig.3C-G). A Med19 protein deleted for its evolutionarily-conserved CORE domain (ΔCORE) still bound Pnr 1-291 (Fig.3D). Binding was also retained after truncating both C-ter and HIM domains, but was abolished if the deletion included the C-terminal end of the CORE domain (aa 126 to 165) (Fig.3E). Deletions starting from the Med19 N-terminus indicated that a truncated protein containing HIM and Cter domains also interacts with Pnr 1-291 (Fig.3F). Further deletions revealed that one fragment of HIM from aa 206 to 220 was critical for Pnr binding in the absence of the CORE domain. Taken together, our data suggest the presence of two Pnr binding sites within Med19, aa 126 to 165 of the CORE (BS1) and aa 206 to 220 of the HIM domain (BS2) (Fig 3C). To further assess their implication, we deleted both BS1 and BS2 in an otherwise full-length Med19 protein. As shown in Fig.3G, the ΔBS1-BS2 mutant no longer bound GATA/Pnr, indicating that at least one of the two binding sites is necessary.

In conclusion, our *in vitro* binding assays indicate that the GATA/Pnr C-zinc finger domain, including its basic tail, binds two separate domains within the evolutionarily-conserved Med19 CORE and HIM regions. Whereas Med19 appears only to bind the C-ZF, Med1 has been shown to bind both GATA/Pnr C-ZF or N-ZF domains *in vitro* (32).

### Med19 physically interacts with GATA/Srp

To investigate whether Med19 is a general GATA cofactor, we asked whether it is able to interact with Serpent (Srp), another *Drosophila* GATA TF family member, as it does with Pnr. First, we used similar GST-pull-down assays (Fig.4A). They showed that recombinant GST-Med19 protein bound *in vitro*-translated full-length GATA/Srp protein. As previously shown for GATA/Pnr, when assaying Srp truncated forms, binding was only retained with the ZF-containing middle part. Splitting the Srp protein in two halves and separating both zinc finger domains indicated that only the C-ZF one is involved in binding Med19 (Fig 4A), as it is also the case for GATA/Pnr (Fig 3E).

**Fig. 4:**
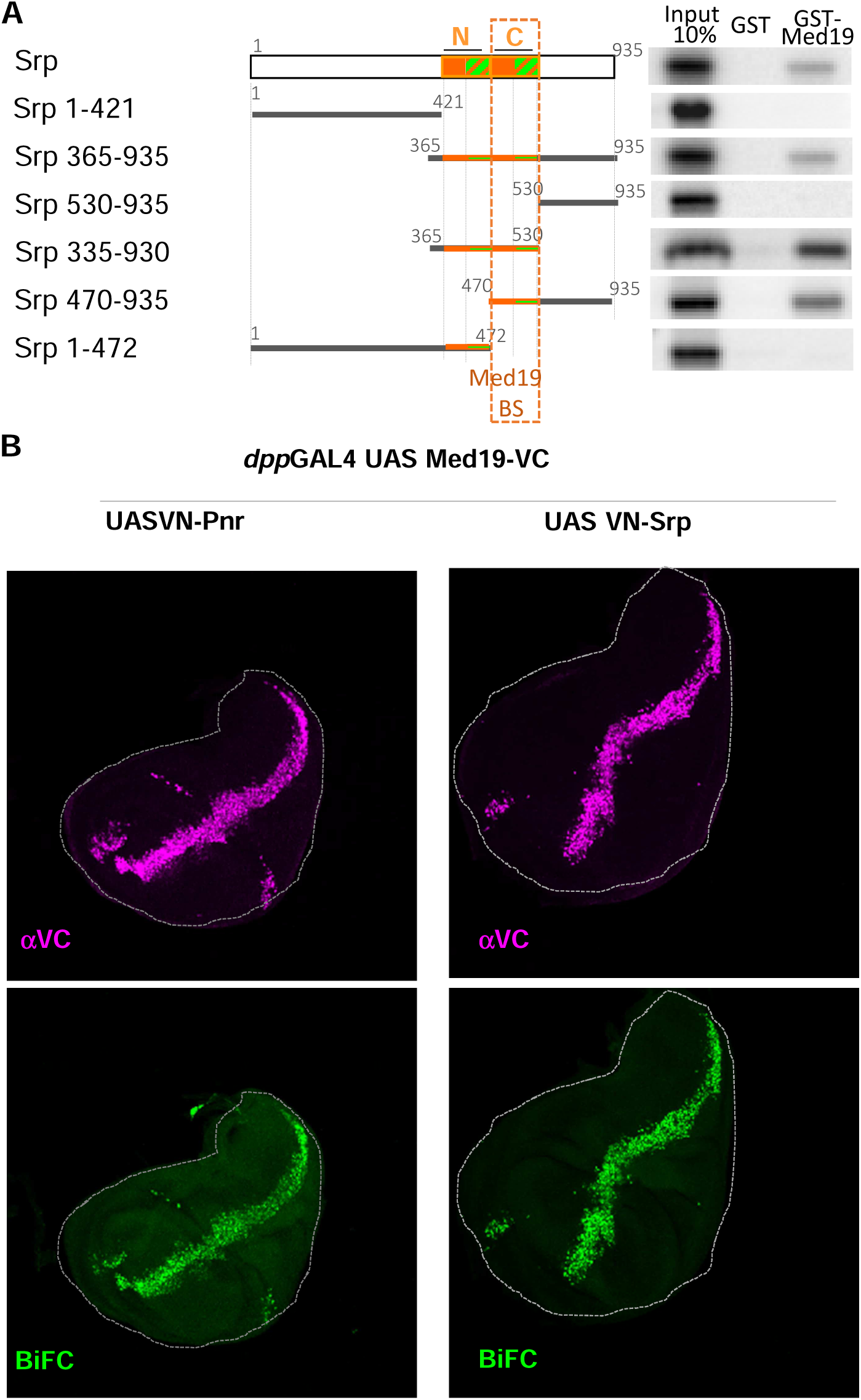
Med19 physically interacts with GATA/Srp. Schematic representation of GATA/Srp and the multiple Srp fragments generated to probe for binding to full-length GST-Med19 (**A**). Autoradiographs from GST pulldown experiments are shown on the right side for each fragment. Again, the critical binding domain is restricted to the GATA C-ZF domain containing the zinc-finger proper (orange square) and its basic tail (hatched green and orange boxes). (**B**) Bimolecular Fluorescence Complementation (BiFC) assays using the *dpp*GAL4 driver to express Med19-VC together with VN-Pnr or VN-Srp. Immunostaining shows similar expression of VC constructs (pink).

To test whether this Med19-Srp interaction also occurs *in vivo*, we used again the BiFC experimental approach. Upon overexpression of Med19-VC with VN-tagged Srp in the *dpp* expression domain, we observed a strong BiFC signal, similar to what we obtained with Pnr-VN (Fig. 4B), indicating that Med19 and GATA Srp indeed interact *in vivo*.

Altogether, these results show that, Med19 interacts both with Pnr and Srp, *in vivo* and, *in vitro* via the GATA family-defining C-zinc finger domain, indicating that Med19 is a general GATA cofactor.

### Med19 shares GATA-cofactor functions with the Med1 Mediator subunit

We previously showed that Med1, another subunit of the MED middle module, is required for Pnr and Srp TF activities *in vivo* and interacts directly with Srp and Pnr, in this case through both their N- and C-zinc finger domains (32). Our new data showing that Med19 can also act as GATA cofactor thus raises the question: do Med1 and Med19 regulate the exact same GATA target genes or do they play distinct roles? Respective to Pnr-dependent transcriptional activity, we have shown that like Med1, Med19 is cell autonomously required for DC-*ac*-lacZ reporter expression whereas *wg* expression requires Med19, but not Med1. This prompted us to consider each GATA target gene as a particular case that could involve interaction with different MED subunits. Kuuluvainen and collaborators (38) identified a set of Srp target genes in *Drosophila* S2 cells, which could be used to test the impact of *Med1*- or *Med19* mRNA depletion. Here, we quantified the expression of 6 Srp target genes: *SrCl, CG14629, CG8157, arg*, and *CG34417 which* are activated (positive targets), and *CG13252* which is repressed by Srp, using Real Time quantitative PCR (qRT-PCR) on control, *Med1*- or *Med19-*mRNA depleted S2 cells. Quantification of mRNAs coding for the Myosin light chain (*Mlc2*) served as a control for housekeeping transcription. As shown in Fig. 5A, in cells depleted for 85% of *Med19* mRNA, expression of the 5 activated Srp target genes *SrCl, CG14629, CG8157, arg and CG34417* was significantly down-regulated and the Srp-repressed target gene *CG13252* was instead up regulated, indicating that Med19 is required *in cellulo* for GATA/Srp transcriptional activity. In cells depleted for about 75% of the *Med1* transcript (Fig. 5B), expression of Srp target genes followed the same trend than after *Med19* mRNA depletion, although less efficiently.

**Fig. 5:**
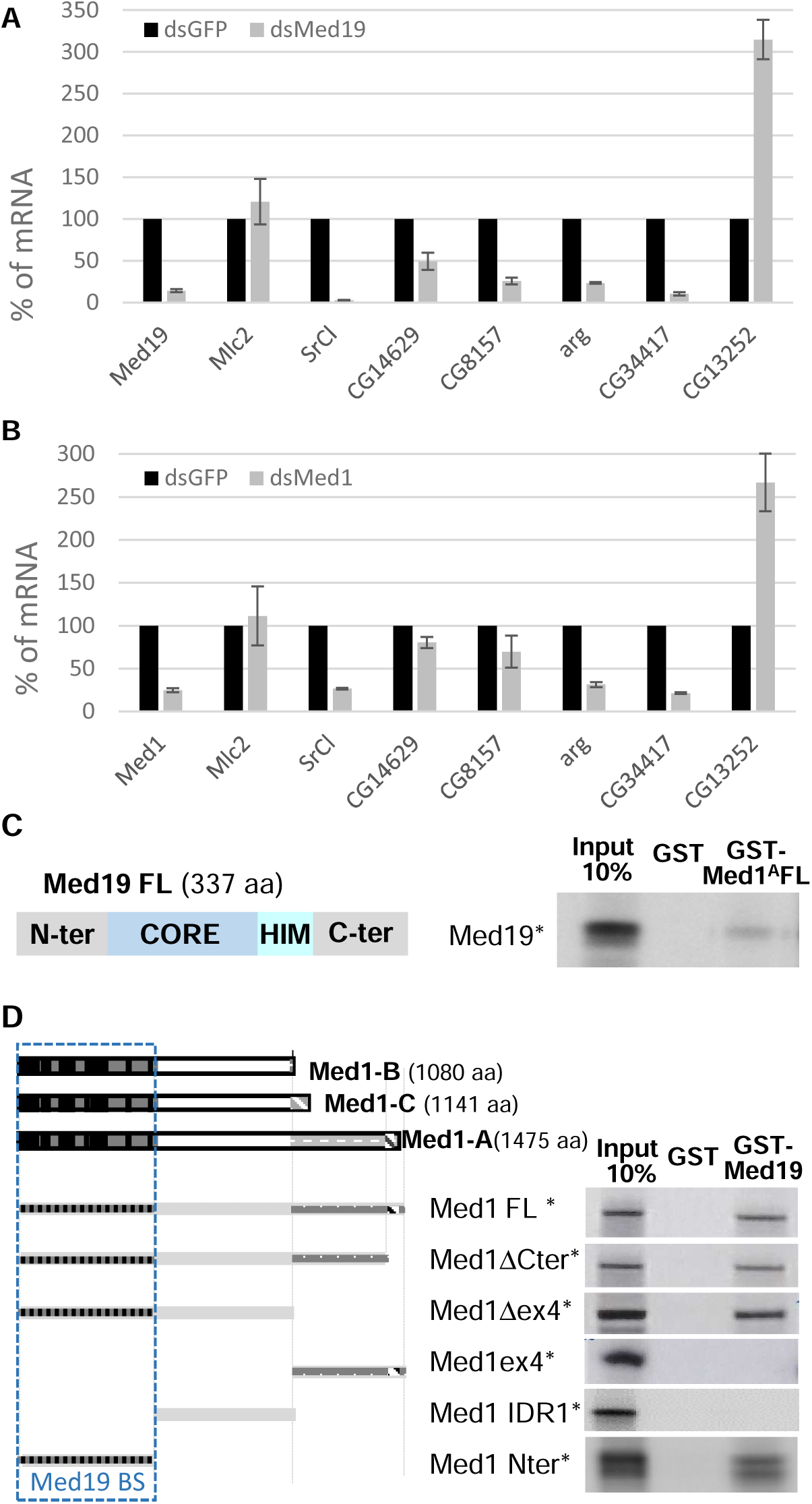
Med19 and Med1 are both required for GATA/Srp target genes expression and interact together. Real Time quantitative PCR analysis of mRNA expression in control cultured cells (dsGFP, **A** and **B**), Med1-depleted cells (dsMed1, A), or Med19-depleted cells (dsMed19, B). The experiment was reproduced 3 times and with a second dsRNA against Med1 and Med19. Autoradiograph from GST pulldown experiments between GST-Med1^A^ and full length Med19 reveal direct interaction between the two MED subunits (**C**). (**D**) Schematic representation of the three Med1 protein isoforms (Med1-A, -B and -C) and the multiple Med1 fragments generated to probe for binding to full-length GST-Med19 are schematized. The N-ter darker grey rectangles correspond essentially to the yeast Med1 orthologue and comprise 15 short evolutionarily-conserved motifs (black boxes (36)). The white and C-ter light grey regions emphasizes the divergent long metazoan-specific extensions. Autoradiograph are shown on the right side, and narrow the Med19-binding domain to the highly conserved N-terminal portion of Med1 proteins.

Several conclusions can be drawn from these experiments: (1) they show that contrary to other MED components, Med1 and Med19 are not generally required for PolII-dependent transcription given that some genes are unchanged or even upregulated. (2) Med19 and Med1 are both required for Srp-mediated gene regulation in cultured cells, seemingly on the same target genes, both for activation and repression (although it is not known whether CG13252 repression by Srp is direct or not).

### Med19 and Med1 can interact directly

Given the shared functional implication of both Mediator subunits in mediating GATA activity, we lastly asked whether Med1 and Med19 proteins are able to interact physically using GST pulldown assays. As shown in Fig.5C, we observed that a GST fusion of Med1 largest isoform A (Med1^A^) bound *in vitro*-translated full-length Med19. In the reverse experiment, we showed that purified GST-Med19 also bound *in vitro* translated Med1^A^ (Fig.5D), showing that both proteins indeed bind to each other *in vitro*, in absence of other MED subunits.

We next sought to identify Med19–interacting domain(s) within the Med1 isoforms by analysing truncated proteins. As shown in Fig.5D, the Med19 interacting domain lies within the evolutionarily-conserved Med1 N-terminal part, which has been proposed to be required for its incorporation within the MED complex (39) and is shared by the three Med1 isoforms. Conversely, Med1 isoform-specific parts are not required for Med19 interaction.

Taken together, GST-pulldown data reveal a direct interaction between Med1 and Med19 which could not be anticipated from structural data (see Fig.6) given that Med1 and Med19 are supposed to lie in opposite parts of the Mediator complex middle module (40).

## Discussion

In this work, we establish using molecular, cellular and genetic analyses that *Drosophila* GATA factors’ transcriptional activity depends on the activity of two Mediator complex subunits, Med19 and Med1. Whereas Med1 could bind both GATA zinc-finger domains (32), we show that Med19 only interacts with the C-ZF domain which also serves as the GATA DNA-binding domain (Fig. 6A). Our analysis of both Pnr and Srp target gene expression in Med19- or Med1-depleted cells indicate that both subunits are critical for multiple GATA target genes regulation, suggesting a close cooperation of Med19 and Med1 to mediate GATA-driven signals. However, at least the Pnr target *wg* relies on Med19 but not Med1, suggesting that the use of MED SU by GATAs can vary depending on the target gene.

**Figure 6:**
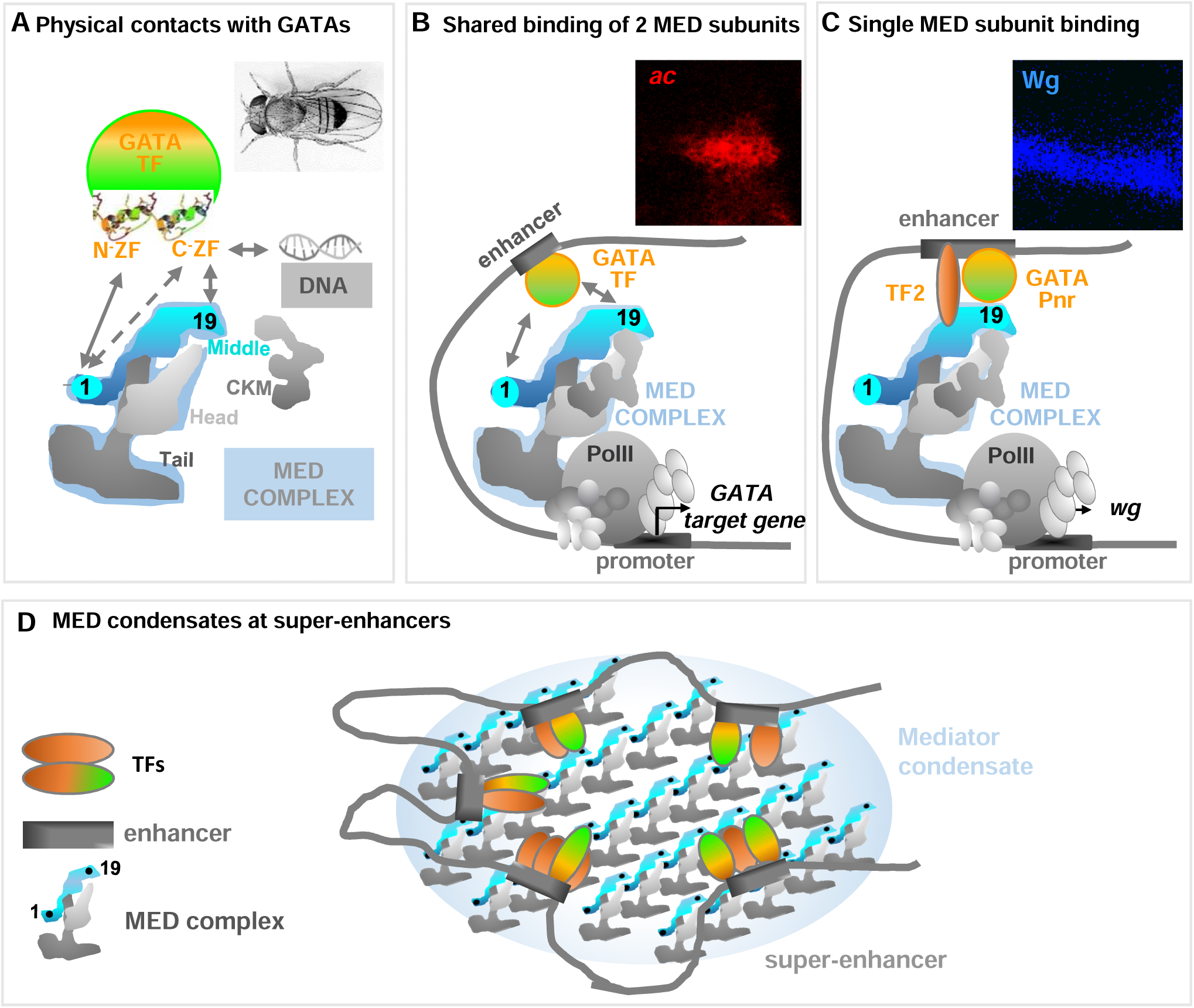
Models of MED subunits – GATA TF interaction.

### New models for GATA – MED interactions

Several models can be envisioned respective to Med19 and Med1 regulatory activities. In the first one (Fig. 6B), we propose that enhancer-bound GATAs use their evolutionarily-conserved ZF-containing domain to directly contact not only the previously identified cofactor Med1 but also the Med19 subunit to recruit the Mediator complex and thus the PolII machinery at some GATA target genes (such as the Pnr target gene *ac*), both contacts being absolutely required. In some cases such as wg (Fig. 6C), Med19 is critical but Med1 is dispensable, so we propose that the presence of other TFs might help recruiting the MED via other subunits.

Previous models of core MED structure-function analysis suggest that the middle and head modules contact the PolII enzyme and associated general transcription factors (GTFs) while the tail module interacts with sequence-specific TFs (2, 3). Our data show that two different MED subunits, Med1 and Med19, are able to bind GATA factors and required for their function and both belong to the middle module. They emphasize that MED should be viewed as a much more complex interface using diverse MED module subunit combinations to contact different TF combinations thus mediating specific transcriptional responses.

### Evolutionarily-conserved GATA-coactivator functions of Med19?

While Med1 is a known GATA cofactor both in mammals and in *Drosophila* (19, 32), the role of Med19 in mediating GATA transcription regulatory properties had never been investigated until now. Here we show that *Drosophila* Med19 binds GATA factors, via motifs lying within the evolutionary-conserved Med19 CORE and HIM domains. Both of these domains bind to the C-ZF domain of GATAs, which is a hallmark of GATA TF family suggesting that interaction with Med19 is likely to be conserved in mammals. Yet, Med1 depletion experiments in mammalian cultured cells induces defects in only a subset of GATA1-activated genes and does not prevent GATA1-dependent repression (41, 42). Furthermore, in studies of the different blood cell types produced by conditional *Med1* knock-out mice, Med1 has been shown to be critical for erythroid lineages which depend upon GATA1-function but is dispensable for hematopoietic stem cell production and T-cell development which require GATA2 and GATA3, respectively (43). Thus, despite being capable of binding all GATA factors *in vitro*, Med1 is not critical for all GATA functions, which suggests that (an) other MED subunit(s) also bind(s) GATAs to relay their regulatory signals to the PolII machinery. Considering the evolutionary-conservation of interaction motifs within both GATAs and Med19 (this work), we argue that Med19 is a strong candidate as a GATA cofactor in mammals.

### Overlapping DNA-Binding and Activation domains of GATA TFs

TFs minimally contain two domains: the DNA binding domains (DBD), which have been extensively studied and allowed to define different TF families, and transcriptional activation domains (TAD), which often link TFs to the RNA polymerase II machinery, and whose structure and characteristics are less well defined. GATA TFs are characterised by the presence of two ZFs which where, so far, thought to play distinct roles. While the C-ZF appeared to be dedicated to DNA binding, co-activators such as dLMO (44) and FOG (45) were shown to bind the N-ZF. Another Pnr interactor, Chip binds to the C-terminal αhelices H1 and H2 (46). Our present data show that Med19 interacts specifically with the Pnr C-ZF. Full interaction requires both the zinc finger proper with the four cysteines responsible for the “finger” structure and the conserved basic tail which contributes to DNA binding (35, 36). It is the first evidence that the *Drosophila* GATA C-ZF may play a dual role, in DNA binding and as an interface with MED subunit(s). Interestingly, the analysis of GATA ZF evolutionary conservation indicates that N- and C-ZF domains comes from a duplication event and that only C-ZF with its basic tail has been conserved in plants, nematode, some insects and echinoderms. Thus, this transactivation function of GATAs’DBD might represent an ancestral GATA function allowing minimal primitive GATAs, essentially composed of the DBD, to connect the MED complex and thus recruit the transcriptional machinery to regulate its target genes.

It seems that this TF property is not restricted to GATA factors. We have previously shown that the Med19 Mediator complex subunit acts *in vivo* as a HOX coactivator and binds through direct contacts with their evolutionarily-conserved helix-turn-helix homeodomain serving as sequence-specific DNA-binding domain (16). These data also corroborate results from a recent high-throughput approach, looking for trans-activation domains of *Drosophila* transcription factors. This work shows that trans-activation domains of several zinc-finger- (ZF-) and basic Helix-Loop-Helix- (bHLH-) TFs overlap structured DNA-binding domains (47). Altogether, these results identify a novel class of TF characterized by overlapping activation and DNA-binding domains and suggest an emerging Med19 property as a dedicated cofactor directly connecting these TFs DNA-binding domains to the general PolII transcriptional machinery.

### Unexpected direct interaction between Med1 and Med19 middle module subunits

Recently, owing to improvement of electronic microscopy and mass spectrometry techniques, important advances have been made in understanding Mediator 3D architecture. It provides a rather precise structural view of MED subunits’ respective position within the yeast and the mammalian complexes (4, 48– 50). Although not yet performed for any insect MED complex, *Drosophila* MED spatial organization can be modelled given the strong structural similarity observed between yeast and human complexes and primary sequence conservation among animals (37). These data indicate that *Drosophila* Med1 and Med19 are most likely situated at two opposite ends of the middle module, Med1 near the tail module and Med19 within the so called “hook” domain proposed to anchor the separable CKM MED module (Fig 6.A).

How, then, reconcile the proposed MED architecture with our results showing a direct interaction between Med1 and Med19 subunits *in vitro*? All the more so as our data suggest an evolutionary conservation of this interaction given that *Drosophila* Med1 binds Med19 through its highly-conserved, N-terminal, MED-addressing domain which is ancestrally present in yeast, as opposed to the C-terminal extensions which are essentially disorganized and fast evolving.

We propose two non-exclusive hypotheses: First, MED complexes could adopt different conformations, which would differ from the “canonical” architecture of the MED complex in isolation. This is supported by observations that the MED complex changes its overall shape when engaged in interactions with either TF, CKM or PolII (49). Perhaps when MED is recruited by GATA, Med1 - Med19 contacts within the MED complex could stabilize one of these “alternative” conformations.

Second, one could envisage that Med1-Med19 interactions do not occur within but between MED complexes and could thus stabilize “multi-MED” structures. It has been shown that master TFs control gene expression programs by establishing clusters of enhancers called super-enhancers, at genes with prominent roles in cell identity (51). Recent studies have revealed that, at super-enhancers, master TFs and the Mediator coactivator form phase-separated condensates, which compartmentalize and concentrate the PolII machinery to specific nuclear foci, to ensure high level of transcription (52–54). Interestingly, mammalian Med1 can form such phase-separated droplets that concentrate the transcription machinery at super-enhancers (Sabari et al., 2018). Bringing together several MED complexes associated with TFs via Med1-Med19 trans-interaction might thus help phase droplets formation at clustered gene enhancers and ensure high transcriptional level (Fig 6.D). Given the more drastic effects of Med19-compared to Med1 depletion on GATA target genes, one could imagine a model where Med19 is absolutely required to recruit MED complex at GATA-bound regulatory sequences whereas Med1 would be preferentially involved in boosting the transcriptional response by favouring the formation of phase-separated MED complex condensates.

In conclusion, our work shows that 2 MED subunits physically bind GATAs and are required to relay the regulatory signals from common TFs. This argues against the generally admitted view of binary interactions between one MED subunit and one (class of) TFs, which appears as an oversimplified model for MED action. The Mediator should be viewed as a complex interface allowing fine-tuned gene regulation by TFs through specific contacts with different MED subunit combinations. This study highlights the unexpected role of *Drosophila* Med19 as a GATA cofactor and Med1 interactor. This work sheds new light on the GATA-MED paradigm and suggests novel means by which several MED subunits might collaborate to regulate gene transcription.

## Experimental procedures

### *Drosophila* stocks, genetic mosaics and phenotypic analyses

Stocks and crosses were maintained at 25°C or 22°C on standard yeast-agar-cornmeal medium. Mitotic clones were generated using the Flp-FRT system with the *Med1*9^2^null allele FRT80B chromosome (16). The Flp recombinase was expressed in the dorsal part of the wing using the *ap*-GAL4 driver recombined with UAS-Flp. The following stocks were used: apGAL4 UASFlp/Cyo; *Med19*^*2*^ FRT2A/TM6B, UbGFP M FRT2A/TM6B; apGAL4^#MD544^ UASGFP/CyO, UAS dsRNA Med19 ^#27559^ and dppGAL4 from BDSC, UAS VN-AbdA and UAS VN-Srp kindly provided by S. Merabet, DC-ac-lacZ kindly provided by P. Heitzler UAS-Med19VC (16) and UAS-Med1^A^VC (32).

Adult phenotypes were analysed by scanning electron microscopy (Hitachi TM-1000 Tabletop model) of frozen adults.

### BiFC assay

This technique is based on expressing *in vivo* two candidate partner proteins with the N- and C-terminal portions (VN and VC) of the Venus protein in order to test the reconstitution of a functional fluorescent protein. UAS-Pnr-VN, UAS-VN-Pnr, and UAS CycC-VC lines were generated by inserting Pnr^A^ or CycC ORF in phase with VN173 (aa 1-172) or VC155 (aa 155-238) ORF in a recipient pUAST-attB plasmid, allowing site specific insertion. attP-carrying embryos expressing PhiC31 integrase in the presumptive germline were injected with these plasmids: Pnr constructions were inserted on the X chromosome (attP ZH-2A), and CycC constructions on the 2^nd^ chromosome (attP 51D) to ensure identical expression of the different VN lines, and easy combination of VN and VC lines. Note that the BiFC constructions were functionally validated for their ability to rescue mutant lethality for Med19-VC, Med1-VC and CycC-VC or to produce typical gain-of-function phenotypes for VN-Pnr.

Crosses were carried out at 22°C for interaction tests to express candidate proteins at homogenous and relatively low levels in order to avoid non-specific signal. We used the *dpp*GAL4 driver (Gal4/UAS system) to direct co-expression of fusion proteins at the antero-posterior frontier of the wing imaginal disc both in the thorax where *pnr* is normally expressed and in the wing pouch region that does not express *pnr*.

### Quantitative analysis of BiFC

Image acquisition was performed on a Leica SP5 using the same settings and number of z slices for the different genetic contexts. BiFC fluorescence was quantified using ImageJ software. Region of interest (ROI) was hand-drawn following the contours of VC tagged protein expression as visualized by antiVC staining. The pixel value sum in the ROI of the BiFC channel was used for comparative quantification. The mean value was calculated from at least 20 wing discs from wandering L3 larvae of each genotype. In each disc, the same area located in the adjacent wing disc tissue was used to estimate tissue auto-fluorescence and remove this noise from the quantification.

### Co-immunoprecipitation experiments

Cultured *Drosophila* S2 cells were grown in 10% serum containing Schneider’s medium at 25°C, and transfected using FuGENE HD transfection reagent (Roche) following manufacturer recommendations. Transfections of 18 x 10^6^ cells per plate were carried out with *pActin*-Myc-Pnr (3µg) and p*Actin*-GAL4 (2µg) plasmids. The cell harvest, protein extraction and IP were performed as described in (32) with the following modifications: the buffer used for protein extraction and subsequent IP contained 0,1% NP40 instead of 0,5% to increase purification of large complexes such as MED; 1mg of total protein extract was used for each IP instead of 1,5mg. Anti-Med19 and non-relevant (NR) IPs were performed with 5µl of decomplemented serum from a Med19-immunized guinea pig (16) or 5µl of the same animal’s pre-immune serum, respectively. We used 10µl G-protein coupled Sepharose beads per IP (SIGMA, P3296). Anti-Myc IPs were performed with 10µl Anti-Myc-Agarose bead (SIGMA, A7470).

Med19 was revealed using a polyclonal serum from guinea-pig (diluted 1:500); Myc-Pnr, with rabbit anti-Pnr (kind gift from G. Morata, diluted 1:1000). We used Lumi-Light^PLUS^ western blotting substrate (ROCHE, 12015196001) and high performance chemiluminescence film (Amersham Hyperfilm™ ECL, GE healthcare, 28906837) for revelation.

### GST-pulldown experiments

Preparation of GST fusion proteins, ^35^S-Methionine-labeled proteins and pulldown were performed essentially as described in (Mojica et al 2017). Med1, Pannier and Serpent proteins or sub-fragments have been produced from cDNA corresponding to Pnr^A^ (Ramain et al 1993), Srp^B^ (Waltzer et al 2002) and Med1^A^ (Immarigeon et al 2019) by in vitro transcription/translation coupled reactions using rabbit reticulocyte extracts (TnT-Promega) isoforms labeled. cDNA encoding full-length dMed19 and deletion derivatives were amplified by PCR using appropriated oligonucleotides and inserted into the *Bam*HI/*Not*I site of the pGEX-6P1 vector (GE Healthcare). Pnr aa 1-291 fragment-encoding was amplified by PCR and cloned into pcDNA3 vector, with an HA-tag at the C-terminus and a Flag-tag at the N-terminus. All clones were verified by sequencing. Primers sequences and complete clone sequences are available upon request. Bacterial expression vectors pGEX-6P1 were transformed in competent E. coli strain BL21 (DE3). The transformed cells were plated in LB agar containing 50 µg/ml of ampicillin. A single colony was grown overnight in 25 ml of LB medium containing ampicillin on a rotary shaker (180 rpm) at 37°C. Overnight starter culture was diluted 1:30 and bacteria were grown in 150 mL of LB medium containing ampicillin at 37°C to an optical density of 0.8-0.9 at 600 nm and expression was induced with 0.5 mM IPTG for 2 h at 37°C. Bacteria were pelleted by centrifugation and pellets were stored overnight at - 20°C. Pellet was resuspended in 15 ml lysis buffer (50 mM Tris-HCl, pH 8.0, 100 mM NaCl, 10 % w/v glycerol, 0.1 % Nonidet-P40) including one Complete™ EDTA-free protease inhibitor tablet and sonicated on ice. After centrifugation at 10 000 x g 45 minutes at 4°C, the supernatant was mixed 2 h at 4°C on a rotating platform with 2 ml Glutathione Sepharose 4B resin. Beads were washed four times with lysis buffer and stored at 4°C.

6µl of tagged Pnr 1-291-HA *in vitro* translation product was mixed with 50 µl of glutathione-agarose bead-GST Med19 derivatives in 200 µl of pull-down buffer (50 mM Tris-HCl, pH 8.0, 100 mM NaCl, 10 % w/v glycerol, 0.1 % Nonidet-P40, 10 µM ZnSO4). The mixture was incubated for 2 h at 4°C, washed four times with 500 µl pull-down buffer. Protein complexes were eluted from the beads with 2X Laemmli sample buffer, boiled 5 min and separated by SDS-PAGE on Mini-PROTEAN® TGX™ precast gels (Bio-Rad). Bound Pnr 1-291 was identified by Western-blot (1:5000 rabbit anti-HA polyclonal) using an ECL kit (Amersham GE Healthcare Life Sciences) based on the manufacturer’s recommendations. HRP-conjugated secondary antibodies were used at 1:5000 and were purchased from Amersham GE Healthcare Life Sciences. All the membranes were scanned on an ImageQuant LAS 500 (GE Healthcare Life Sciences).

### Sequence conservation analysis

C-ZF domains from the *Drosophila* and human GATA family members were extracted from the NCBI web site (https://www.ncbi.nlm.nih.gov/protein/), aligned with the MAFFT software (https://mafft.cbrc.jp/alignment/server/), using default parameters, and amino acid conservation were visualized with Jalview 2.10.5 version (http://www.jalview.org/) using Clustal coloring.

### RT-qPCR

Two different dsRNA were used for *Med1*- or *Med19* mRNA depletion, only one for control GFP mRNA. The indicated dsRNAs (see Table I is added at 2µg/ml to exponentially growing S2 cells, in an orbital shaker, at 2 10^6^ cell/ml in serum-free medium. After 40min, serum is added. 24h later a second addition of dsRNA is done at 1µg/ml. Cells are collected 5 days after the first dsRNA treatment.

**Table 1:**
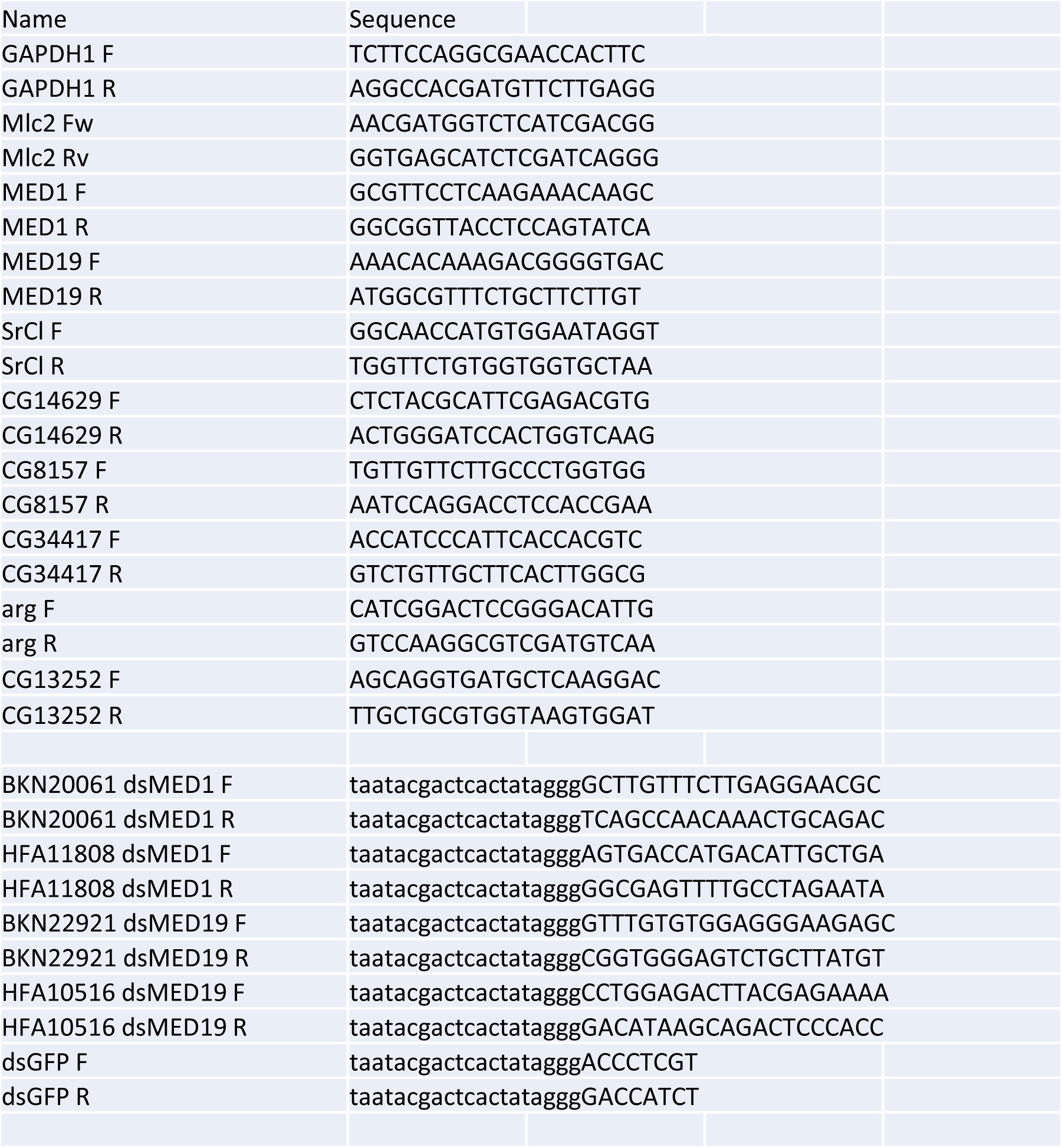
dsRNA used for GFP, Med19 and Med1 depletion in Drosophila cultured cells.

For mRNA quantification, mRNAs were purified by RNeasy kit (Qiagen). Reverse transcription was done using SuperScript™ II Reverse Transcriptase (Thermo Fisher Scientific) and cDNA were quantified by real-Time qPCR (CFX Bio-Rad) with specific oligonucleotides (Table I). Absolute quantification of each mRNA was normalized to *GAPDH* mRNA quantity in the same sample. mRNA measured in cells treated with a control dsRNA GFP was set at 100% to compare with cells treated with a dsRNA against Med1 or Med19.

## Acknowledgements

We gratefully acknowledge P. Heitzler, S. Merabet and BDSC for plasmids and fly stocks, D. Jullien, A. Vincent and C. Monod for critical reading of the manuscript, V. Gobert and B. Augé for technical help and finally the Toulouse Rio Imaging platform.

## Fundings and additional information

This work was supported by grants from the French “Ministère de l’Enseignement et de la Recherche” (C.I.), the Ligue Nationale contre le Cancer (C.I. and A.P.), the Association pour la Recherche sur le Cancer (ARC PJA 20141201932), the Agence Nationale de Recherche (ANR-16 CE12-0021-01), and we benefitted from the support of the Centre National de Recherche Scientifique (CNRS) and Toulouse III University.

## Supporting information

**Figure S1:**
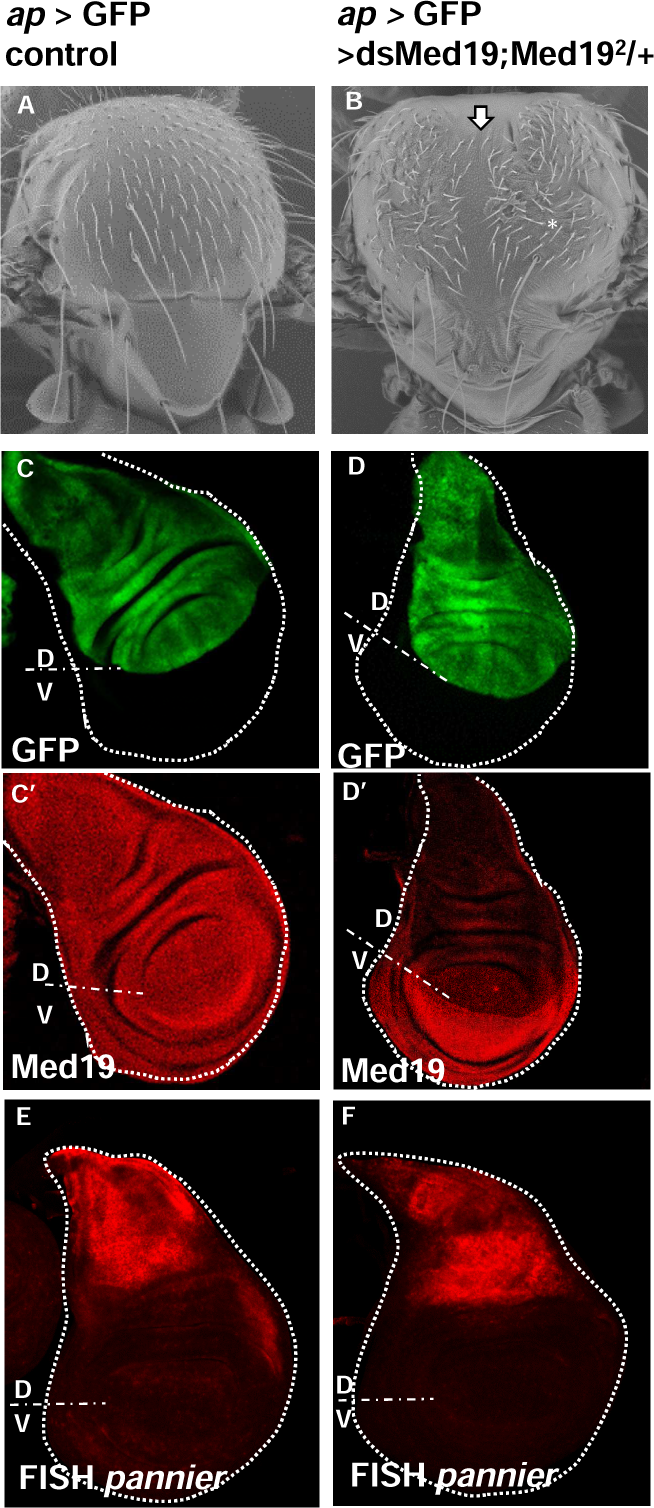
Med19 depletion produces adult thoracic cleft and DC macrochaete loss, but does not change larval pnr expression

